# Longitudinal changes in control energy of brain networks in older adults at familial risk for Alzheimer’s disease

**DOI:** 10.64898/2026.09.06.749740

**Authors:** Paulina Skolasinska, Alfie Wearn, Daniel O’Sullivan, Isabella Stallworthy, Julia K. Brynildsen, Ilana R. Leppert, Christine L. Tardif, Gary R. Turner, Dani S. Bassett, R. Nathan Spreng, the PREVENT-AD Research Group

## Abstract

The ability of structural brain networks to control neural dynamics is affected by healthy and pathological aging, including Alzheimer’s disease (AD). According to network control theory, transitions between functional brain states incur energetic costs. How these costs change longitudinally with aging remains unknown. Here, in 279 older adults at familial risk for AD, control energy increased longitudinally for state maintenance and transitions between sensorimotor and attentional networks, while decreasing for another sensorimotor–attentional transition. Changes in transition costs between higher-order networks were associated with cognitive performance: greater DorsAttn⟷Executive cost was associated with poorer immediate memory, while lower Executive⟷Def+Lim cost was associated with poorer attention. Higher plasma p-tau181 was associated with lower DorsAttn⟷Def+Lim transition cost. These findings reveal transition-specific alterations in the energetic landscape of aging and distinct relationships with cognition and AD pathology. Together, they highlight energetic constraints on brain network reconfiguration as a complementary perspective on neurocognitive aging.

---

Aging is associated with substantial changes in the organization and dynamics of large-scale brain networks, including alterations in functional connectivity, network segregation, and the brain’s ability to flexibly reconfigure in response to changing cognitive demands^1–3^. While functional connectivity and brain state approaches can characterize how network organization and dynamics change with age, they do not account for the structural architecture that constrains these dynamics. Age-related decline in white-matter integrity and structural connectivity^4,5^ together with increasingly inefficient patterns of functional organization may place greater energetic demands on the brain as it strives to maintain and reconfigure between distinct brain states^6^, that is temporary and recurring patterns of synchronized activity. These energetic demands may become pronounced in the context of accumulation of neuropathology and the development of Alzheimer’s disease (AD), further limiting the brain’s ability to dynamically reorganize its activity.

Network control theory (NCT) is a computational modelling approach which offers a complementary framework for quantifying these energetic demands by modeling how the brain’s structural architecture constrains transitions between functional states and estimating the energy required to maintain or reconfigure them. Thus, NCT provides a means to move beyond describing age-related changes in brain dynamics toward understanding their underlying energetic constraints. NCT has been leveraged to understand brain functional dynamics in healthy younger^7–9^ and older adults^10–12^, to better understand disease etiology and pathology in clinical populations (e.g., epilepsy^13,14^ and Parkinson’s disease^15^) including AD^16,17^ to model neural dysfunction, elucidate disease mechanisms, and to inform the design of therapeutic neuromodulation^14,17^. However, applications of NCT to the study of the effects of aging and AD-related pathology on the energetic demands of regulating brain dynamics remain limited.

A sole investigation of age-related differences in energy costs showed differences in the relationship between regional control energy patterns of transitions from sensory to attention networks and working memory performance between older and younger adults^12^. Meanwhile, another study observed lower transition energies of the limbic network in mild cognitive impairment (MCI) patients compared to controls^18^. Although the study did not find a link between transition energies and AD-related neuropathology (amyloid and tau) derived from positron emission tomography (PET) using regression models, node-level information about cortical features can be incorporated into the NCT model itself to assess their impact on the energetic cost of transitions^7–9,19^. This approach could be particularly appropriate for incorporating markers of AD pathology into control analyses.

Overall, the existing literature suggests NCT might be a valuable tool for characterizing the neural mechanisms behind the changing energetic landscape of the aging brain. Despite these insights, none of the studies mentioned above performed gold-standard longitudinal investigations of the energy costs required for transitions between brain states in aging or along the AD spectrum. Further, it remains necessary to probe the relationship between age-associated changes in the energetic landscape and cognitive function across multiple domains, in addition to probing relations with markers of AD pathology. Addressing these gaps, the current study also advances previous NCT methodologies by incorporating additional biologically relevant data. To build the structural connectomes that form the basis of the NCT model, we enriched the tractograms derived from the diffusion weighted images with white matter microstructure data with myelin-sensitive magnetization transfer saturation (MTsat). These enriched structural connectomes provided the anatomical constraints for modeling the propagation of functional activation obtained from multi-echo fMRI.

Hence, our application of NCT integrated state-of-the art multimodal MRI and PET data in a longitudinal sample of 279 older adults at familial risk of AD. Our aim was to characterize the longitudinal changes in the energy costs associated with transitions between brain states as well as maintenance of ongoing states in the context of aging and AD risk. We expected increases in energy costs, or reduced transition efficiency, due to established deterioration of white matter pathways which guide the modeled transitions. Furthermore, we leveraged the densely sampled PREVENT-AD dataset to contextualize the observed changes in terms of AD-related neuropathology and multi-faceted cognitive function. We hypothesized that the increased energetic costs would be accompanied by accumulation of pathological changes and greater cognitive decline. Overall, our work furthers an understanding of healthy and pathological neurocognitive aging through the lens of brain energetics.

## Results

To assess whether aging is related to an increase in the energetic cost of switching between brain states or maintaining states, we applied NCT to multimodal MRI data in a longitudinal cohort of older adults at familial risk for AD (PREVENT-AD; Table 1). To determine whether some brain networks or transitions are particularly vulnerable to age-related increases in energetic cost, we assessed these costs between each pair of network-based brain states. Additional weighted models separately incorporated cortical volume, amyloid-PET, and tau-PET cortical maps to explore how these features may influence the aforementioned brain state energetics (Figure 1). Further, to assess whether the age-related changes in energetic control costs were more heavily influenced by the functional states or the structural architecture, we tested separate models with either individual-level function and group-averaged structure or individual structure and group-averaged function. Finally, to determine the broader biological and behavioral significance of energetic costs, we examined whether they were associated with AD risk and pathology as well as cognitive function across multiple domains.

**Table 1.**
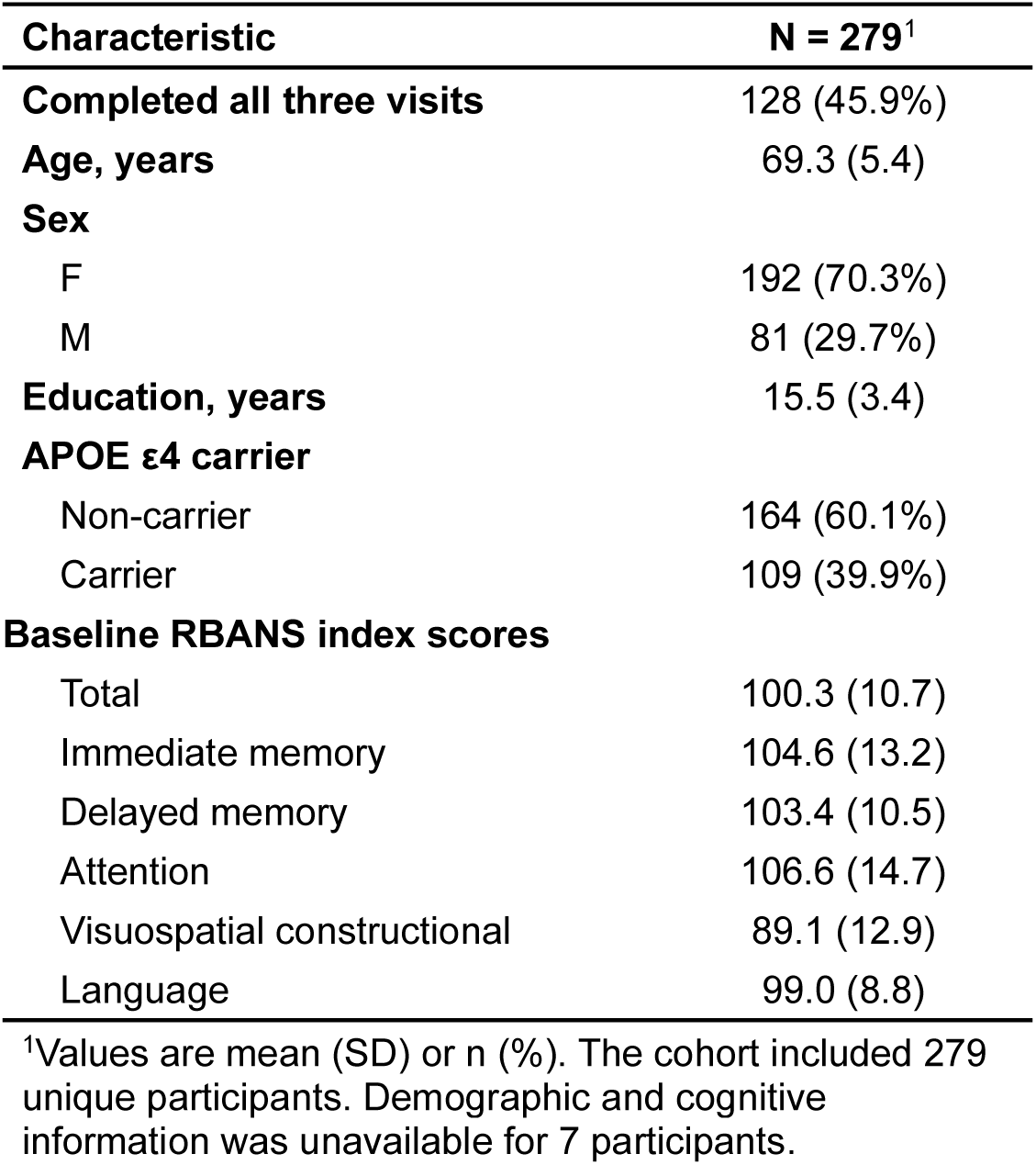
Participant characteristics.

**Figure 1.**
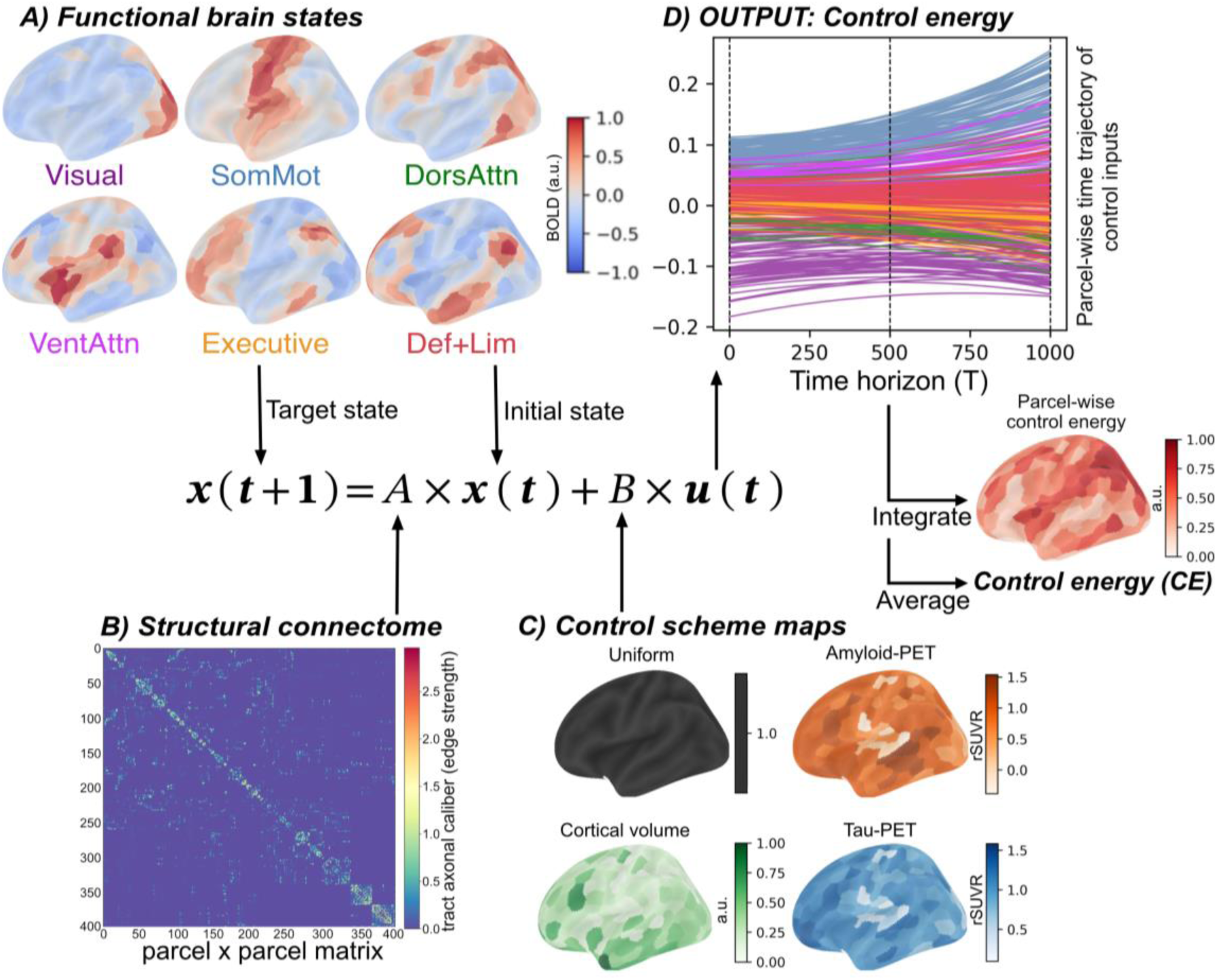
Multimodal neuroimaging data was used to inform a network control theory (NCT) model to obtain the control inputs required to complete a brain state transition. As the dynamic system evolved across a continuous time horizon (*T*), the control inputs (*u*) and brain activations (*x*) were calculated at each time point (a.u) from the initial (*x(t)*) to the target (*x(t+1)*) state following the optimal transition trajectory constrained by the structural connectome (*A*). Multiple control schemes were used so that the control inputs were either uniform or weighted by cortical features (*B*) obtained from structural MRI and PET scans. A) Pairs of brain states corresponding to the canonical Yeo *et al.* 2011 networks (with default and limbic networks combined) were created according to the method described in Caballos *et al*. (2022), from 3T multi-echo functional MRI. Each brain state was used as either the initial or the target state or both, to control inputs associated with between-network state transitions or that related to maintaining a state when initial and target states were the same. B) Microstructure-informed structural connectomes weighed for tract caliber were created by combining diffusion-weighted imaging (DWI) and multiparametric mapping (MPM) that included a sequence weighed for magnetization transfer saturation (MTsat). The activation was guided from the target to the initial state along the edges of the structural network. C) Multiple control schemes were applied in separate models. They included uniform control where each node was weighted by the same value of 1 or control weighted by cortical features such as cortical volume obtained using *freesurfer* from the T1w MRI and cortical amyloid and tau obtained from PET scans in a subset of participants. Processing of the PET images was done according to the method described in Wearn et al. (2026). D) Control energy or energetic cost of each state transition was obtained by integrating the corresponding control inputs in time and averaging across all nodes. Data from all modalities was parcellated into 400 Schaefer parcels.

### Transition costs differ between network pairs at baseline

Due to the novelty of the NCT model, especially in older adult samples, contextualizing findings are scarce. Therefore, before assessing whether energetic costs of state transitions changed in aging, we first examined the characteristics of these costs at baseline. From the NCT models, we extracted the energetic costs of transitions between each pair of functional brain states and averaged the symmetric transitions (i.e., A⟷B and B⟷A). We also extracted the costs of maintaining states (i.e., A⟷A or B⟷B) and averaged them across all network states.

The energetic costs differ based on the networks involved in the transition in younger adults^20^. A lack of such differences in older adults might suggest a dedifferentiation of the energetic landscape in older age. To test this, we compared the energetic costs associated with each state transition (Figure 2A) using linear mixed-effects models (LMM) with individual intercepts, where the energetic cost was predicted by the transition. Overall, the energetic costs differed significantly between the transitions [F(15, 4448) = 520.1, p < 0.001]. The Executive⟷Def+Lim and SomMot⟷VentAttn transitions were least costly whereas transitions between Def+Lim and the two attention networks (DorsAttn, VentAttn) states were most costly. This suggests that the transitions traversed distinct trajectories incurring varying amounts of cost.

**Figure 2.**
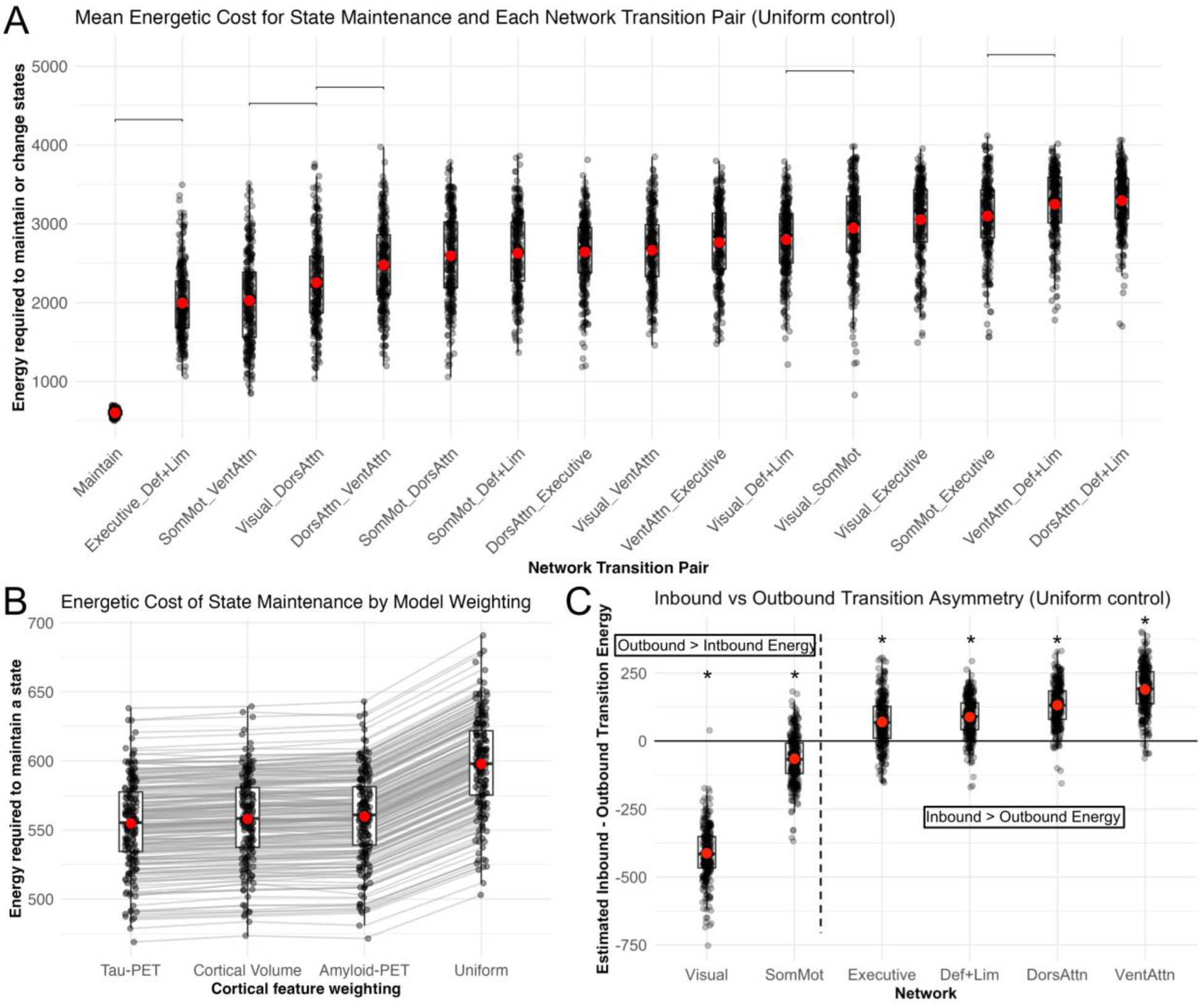
Costs of transitions differed between and within network pairs at baseline. **A)** We estimated and compared the energetic costs related to each pair of state transitions for the unweighted model with the uniform control scheme. Cost of state maintenance was lower than the costs of between-network transitions. Among the latter, Executive⟷Def+Lim and SomMot⟷VentAttn transitions were the least energetically costly while transitions between Def+Lim and the two Attention networks were the most energetically costly. The symmetric transitions (A ⟷ B, B ⟷ A) were averaged. Costs of state maintenance (A ⟷ A) were also averaged across all networks. The brackets were plotted only for the adjacent significant comparisons. **B)** There was a significant difference between the unweighted and weighted models for the costs of state maintenance. Including the information describing cortical features in the NCT model resulted in reduction of energetic costs, with cortical tau providing the greatest cost reduction, from the baseline uniform model, followed by cortical volume and cortical amyloid. **C)** There were significant asymmetries between all inbound and all outbound transitions for each network, so that it was more costly to enter the unimodal states than to leave them, and the reverse was true for the transmodal network states. Further, the strength of asymmetry differed between the networks with the Visual network showing the largest negative asymmetry of transition (inbound > outbound) and the VentAttn network showing the greatest positive asymmetry (outbound > inbound). The brackets and asterisks indicate significant (*p*FDR’s < 0.05) difference from zero after multiple comparison correction.

Previous research indicated spatial distribution of certain features, such as neurotransmitter receptors^8^, affects control inputs in the NCT models. Here, we asked whether imposing biologically informed, spatially heterogeneous constraints on control, in the form of cortical features with high relevance to aging and AD, alters the energetic requirements of brain state transitions. To address this question, we incorporated cortical volume, amyloid-PET, and tau-PET maps into the NCT model by weighting the control-input matrix *B,* thus constraining the availability of control inputs, such that regions with higher feature values have greater influence on the control input than regions with lower values. Greater cortical volume indicated less neurodegeneration, while greater amyloid-PET, and tau-PET values indicated more pathology. We contrasted each weighted model with the non-weighted uniform model for each transition to quantify the contribution of the aforementioned cortical features to control energy (Figure 2B). There was a significant difference between the control energy costs under varying control schemes for all transitions, driven by the large difference between the uniform and the weighted models, so that on average there was a significant reduction of ∼8% in control cost when the model was informed by a cortical feature vs. the non-weighted model. Within the weighted models, the transitions tended to have the lowest cost in the tau-PET model followed by cortical volume and amyloid-PET. The difference between the tau-PET and cortical volume-informed models was small (0.6%) but significant except for the Visual⟷Executive transition. Whereas the difference between cortical volume and amyloid-PET (0.3%) models was significant for state maintenance, transitions involving the visual network, and for transitions between SomMot⟷Executive and VentAttn⟷Executive networks. Significant between-model tests despite small differences in these costs was due to the highly consistent within-subject effect. Overall, the spatial distribution of neurodegeneration and AD pathology was relevant to the energetic demands of brain state reconfigurations, beyond the constraints imposed by the structural connectome alone.

Lastly, we investigated the differences in the transition costs within the symmetric state transition pairs, i.e., transition asymmetry. Our decision to average these symmetric transitions was motivated by their very strong correlation (average *r*=0.97) and the desire to limit the number of comparisons. However, others have found significant differences between transition costs depending on the direction of the transition, that is up or down the cortical hierarchy^19,20^, with the bottom-up transitions being less energetically costly than the top-down transitions^19^. Here, to test whether transition asymmetry is maintained in older age, for each state we calculated the average cost of transitions from any other state into that state, i.e., the inbound energy cost; as well as the average cost of transitions from that state to any other state, i.e., the outbound energy cost. Then, for each network we computed the difference between the inbound and the outbound energy to obtain the transition asymmetry. We found transition significant asymmetries (significant difference from zero; p’s < 0.05) suggesting that the energetic landscape of brain dynamics is directionally structured in older adults. It was energetically easier to enter the unimodal states (Visual, SomMot) than to leave them, indicated by negative transition asymmetry. Whereas transmodal states were energetically costly to enter but relatively inexpensive to leave, indicated by positive asymmetry.

### The energetic costs of state transitions reveal distinct longitudinal change trajectories

Given the deterioration of the structural pathways^4,5^ and declines in the dynamic organization of the functional connectome^1–3^ in aging, we expected to observe longitudinal increases in the energetic costs of state transitions and state maintenance in our cohort of older adults at elevated risk of AD. For this purpose, we fitted LMMs to predict changes in transition costs, modelling each between-network state transition and state maintenance separately. Time since baseline was the main predictor. The models were adjusted for baseline age, sex, years of education, in-scanner motion and estimated intercranial volume. The individual intercept was included.

In the primary uniform model, not weighted for cortical features, there was an increase in the costs of between-state transitions: Visual⟷SomMot (i.e., the two unimodal states) and SomMot⟷VentAttn states. The cost of state maintenance also increased. We also observed a significant decrease in the cost of transitioning between SomMot⟷DorsAttn states (Figure 3). Other state transitions did not change significantly in the study period (Table 2). There were no significant changes in energetic asymmetries (all pFDR’s > 0.05). Therefore, aging was not associated with a uniform increase of transition costs; rather, it there was a selective reorganization of the energetic landscape for particular network interactions, while its directional organization remained unchanged.

**Figure 3.**
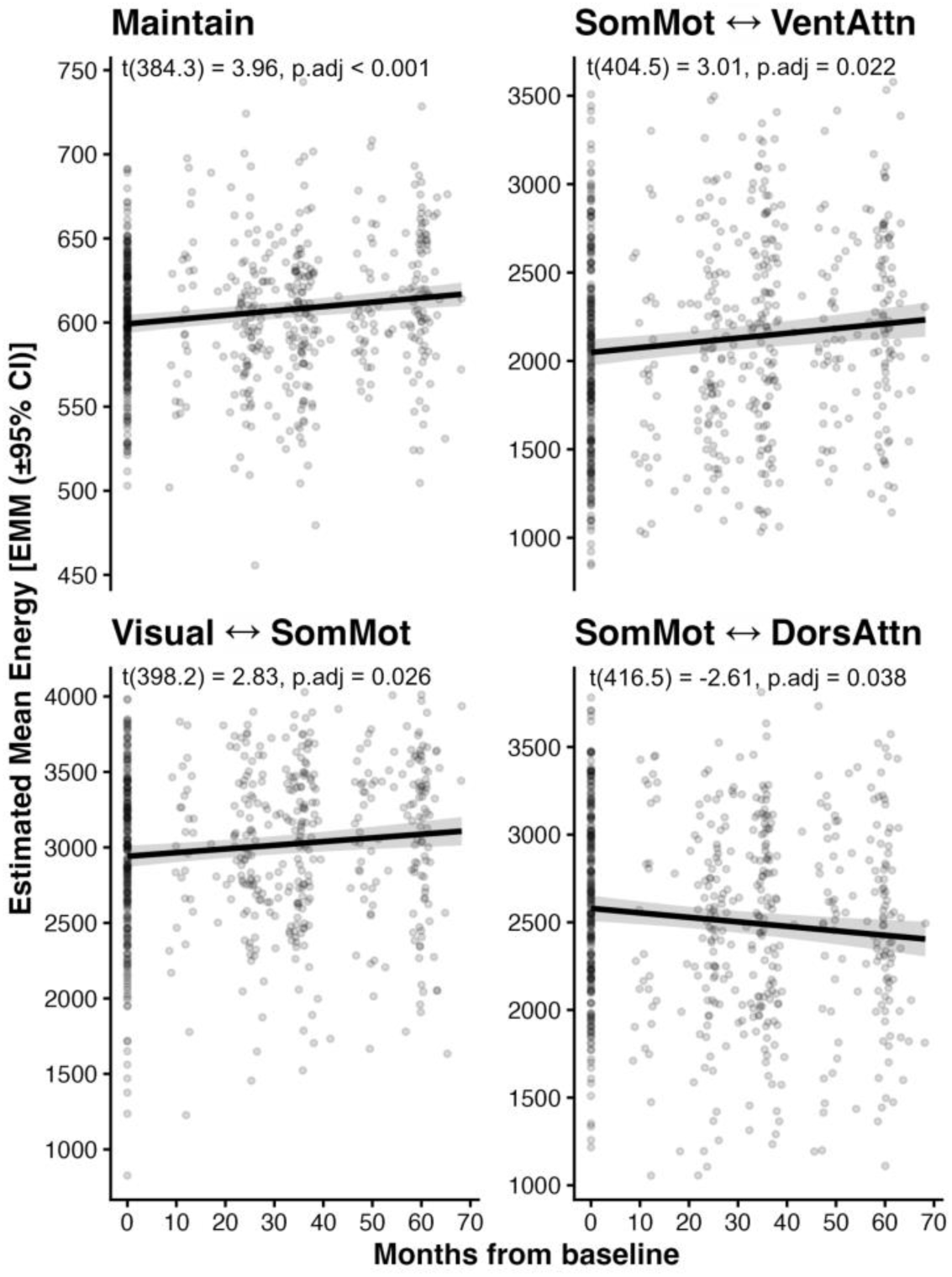
Energetic costs of transitions showed distinct trajectories of longitudinal change. In line with our hypothesis, we observed increased costs for three of the transitions, between the SomMot⟷VentAttn states and between the two unimodal states (Visual ⟷ SomMot), as well as for state maintenance. Conversely, we found a decrease in energetic cost of the SomMot⟷DorsAttn transition. Notably, in the model keeping the functional states constant between the participants (IS/GF model) there was an increase in cost for all transitions pairs. There were no substantial differences between the uniform model and the model weighted for cortical volume, and the models enriched with PET-derived tau and amyloid, except that Visual⟷SomMot transition did not show significant change in the latter two models, likely due to decreased sample size. The p-values were FDR-corrected for 16 comparisons.

**Table 2.** Longitudinal effects of time on transition energy by NCT model (β estimates and FDR-adjusted p-values)

| Transition | Uniform Model | Uniform (Female) | Uniform (Male) | Cortical Volume | Amyloid-PET | Tau-PET | Uniform GS/IF | Uniform IS/GF |
| --- | --- | --- | --- | --- | --- | --- | --- | --- |
| Visual ↔ SomMot | 2.42, 0.026* | 2.29, 0.095 | 2.91, 0.259 | 2.19, 0.032* | 1.70, 0.101 | 1.72, 0.090 | 1.94, 0.043* | 0.11, 0.001*** |
| Visual ↔ DorsAttn | 1.03, 0.398 | 0.24, 0.922 | 3.29, 0.259 | 0.94, 0.404 | 1.24, 0.339 | 1.25, 0.326 | 0.71, 0.526 | 0.19, 0.001*** |
| Visual ↔ VentAttn | -1.66, 0.100 | -2.39, 0.049* | 0.39, 0.984 | -1.53, 0.107 | -1.13, 0.339 | -1.09, 0.326 | -1.55, 0.091 | 0.12, 0.001*** |
| Visual ↔ Executive | -1.54, 0.100 | -1.72, 0.155 | -2.08, 0.477 | -1.39, 0.107 | -1.69, 0.094 | -1.64, 0.090 | -1.42, 0.091 | 0.12, 0.001*** |
| Visual ↔ Def+Lim | -0.15, 0.896 | 0.40, 0.828 | -1.69, 0.641 | -0.14, 0.901 | -0.58, 0.560 | -0.57, 0.560 | -0.25, 0.758 | 0.13, 0.001*** |
| SomMot ↔ DorsAttn | -2.54, 0.038* | -3.09, 0.020* | -1.33, 0.766 | -2.35, 0.035* | -3.11, 0.007** | -3.07, 0.007** | -2.30, 0.043* | 0.18, 0.001*** |
| SomMot ↔ VentAttn | 2.60, 0.022* | 4.58, 0.001*** | -2.19, 0.471 | 2.37, 0.023* | 2.79, 0.007** | 2.76, 0.007** | 2.02, 0.043* | 0.14, 0.001*** |
| SomMot ↔ Executive | 1.38, 0.100 | 1.61, 0.148 | 0.97, 0.753 | 1.25, 0.107 | 1.59, 0.075 | 1.60, 0.071 | 1.06, 0.169 | 0.12, 0.001*** |
| SomMot ↔ Def+Lim | 0.70, 0.420 | 0.14, 0.922 | 2.09, 0.471 | 0.66, 0.404 | 0.70, 0.431 | 0.72, 0.412 | 0.40, 0.600 | 0.17, 0.001*** |
| DorsAttn ↔ VentAttn | 2.33, 0.069 | 3.28, 0.020* | -0.23, 0.996 | 2.13, 0.071 | 2.20, 0.058 | 2.18, 0.057 | 1.82, 0.091 | 0.19, 0.001*** |
| DorsAttn ↔ Executive | 0.11, 0.896 | 1.31, 0.335 | -2.34, 0.366 | 0.09, 0.905 | 0.51, 0.566 | 0.51, 0.563 | -0.06, 0.934 | 0.20, 0.001*** |
| DorsAttn ↔ Def+Lim | -0.76, 0.398 | -0.99, 0.335 | -0.01, 0.996 | -0.68, 0.404 | -0.68, 0.413 | -0.70, 0.389 | -0.79, 0.282 | 0.15, 0.001*** |
| VentAttn ↔ Executive | -0.70, 0.462 | -0.97, 0.477 | 0.08, 0.996 | -0.66, 0.447 | -0.85, 0.431 | -0.85, 0.412 | -0.75, 0.422 | 0.15, 0.001*** |
| VentAttn ↔ Def+Lim | -0.87, 0.343 | -0.07, 0.924 | -2.64, 0.259 | -0.80, 0.334 | -0.84, 0.339 | -0.84, 0.326 | -0.86, 0.251 | 0.12, 0.001*** |
| Executive ↔ Def+Lim | 0.66, 0.398 | 0.70, 0.477 | 0.76, 0.766 | 0.59, 0.404 | 0.43, 0.560 | 0.44, 0.554 | 0.37, 0.600 | 0.22, 0.001*** |
| Maintain | 0.21, 0.001** | 0.29, 0.001*** | 0.03, 0.961 | 0.19, 0.002** | 0.20, 0.002** | 0.20, 0.003** | -0.04, 0.284 | 0.25, 0.001*** |

Values are $\beta$ estimates and FDR-adjusted p-values. \* pFDR < 0.05; \*\* pFDR < 0.01; \*\*\* pFDR < 0.001.
GS/IF = Uniform control with group-averaged structural connectome and individual functional states, IS/GF = Uniform control with group-averaged functional states and individual structural connectomes. All other models used individual structural connectomes and individual functional states. The results disaggregated by sex were investigated in the uniform model.

#### Weighting NCT models by cortical features had limited effect on the change in energetic costs

In addition to the uniform (non-weighted) model, we investigated the longitudinal change in the NCT models where the control inputs (*u*) were weighted by cortical features through the *B* matrix. We hypothesized that incorporating the spatial distribution of cortical volume and AD pathology into the control input matrix would modify age-related trajectories of control energy. We expected cortical volume weighting to attenuate the expected increases in control energy by emphasizing regions with greater preserved cortical integrity, whereas amyloid- and tau-informed weightings were expected to accentuate the age-related increases by emphasizing regions vulnerable to AD pathology. However, weighting the model by cortical volume did not results in substantial differences in longitudinal trajectories compared to those from the uniform model. Similarly, including PET-derived cortical features, amyloid-PET and tau-PET, resulted in the same pattern of results. An exception was the Visual⟷SomMot transition, where the effect of time was non-significant and attenuated, which is contrary to our expectations. However, this might be due to the reduced sample size for the PET-informed models (N=188). Therefore, there was no to little effect of weighting the control schemes on the longitudinal trajectories of control energies.

#### Individual variability in structure and function affect the changes in energetic costs

The primary uniform model, like all weighted models, was estimated using individual-level structural and functional data (IS/IF), thereby capturing subject-specific differences in both structural connectivity and functional brain states. To determine the extent to which individual variation in structural connectivity versus functional brain states contributed to estimates of control energy, we compared the IS/IF model with two additional non-weighted models. The first used group-averaged structural connectivity with individual functional states (GS/IF), while the second used individual structural connectivity with group-averaged functional states (IS/GF). If individual structural connectivity was a primary determinant of control-energy estimates, we expected the IS/IF and IS/GF models to show similar patterns, with greater divergence between IS/IF and GS/IF. Conversely, greater similarity between IS/IF and GS/IF would suggest that individual functional states contribute more strongly to between-person variation in control energy. Comparing these models therefore allowed us to assess the relative contribution of individual structural and functional variation to control energy estimates and to determine whether longitudinal effects were robust to the use of group-averaged structural or functional data.

We found that when the structural connectome (the *A* matrix in the NCT model) was kept constant between the participants, the effects were similar but attenuated compared to the IS/IF model. State maintenance was the exception, as it showed a small negative albeit not significant slope for time, which was positive in the main uniform model. When the states were shared between the participants, but everyone retained their individual structural connectomes (IS/GF model), we observed a universal increase in energetic costs for all between-state transitions and state maintenance. This pattern, together with the marked longitudinal decline in average structural connectivity [t(371) = -13.9, p < .001], suggests that degradation of the structural substrate may impose increasingly costly constraints on brain state control. Table 2 includes the β coefficients and p-values for the time predictor of all tested models.

### Higher level of Alzheimer’s disease pathology was related to facilitated transitions between antagonistic networks

We expected higher levels of AD pathology, p-Tau181, and the presence of AD risk factors to be associated with higher energetic costs of transitions, due to the relationship between negative relationship between p-Tau181^21^, presence of the APOE4 gene^22^, cognitive decline^23^, and white matter integrity.

To assess the effect of AD risk and pathology on energetic costs of transitions and their longitudinal trajectories, we fitted LMMs with random intercepts, with AD pathology (p-Tau181) and AD risk factors (APOE4 carriership status and cognitive vulnerability) in interaction with time since baseline as the main predictor. Each risk factor and pathological marker was tested in a separate model. The energy costs of each between-network state transition and of state maintenance were also modeled separately. The models were adjusted for the same set of covariates as the main set of longitudinal models.

Lower cost of transitions between DorsAttn⟷Def+Lim was related to higher levels of p-Tau181 after FDR correction for 16 comparisons. The main effect p-Tau181 was significant, but not its interaction with time since baseline. There were no associations between energetic costs and p-Tau181 for the other state transitions or for the two AD risk factors (Figure 4A).

**Figure 4.**
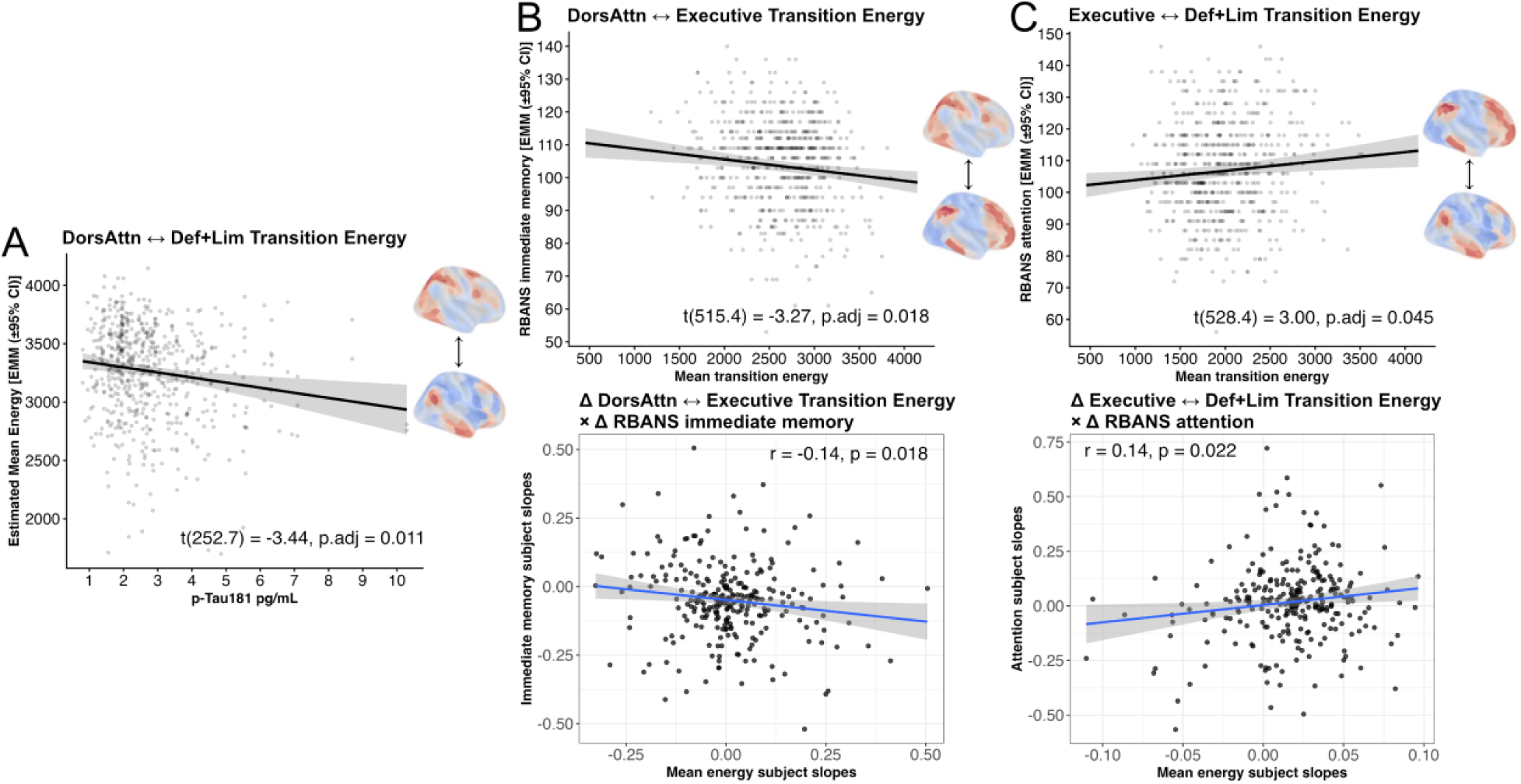
Transition costs between some higher-order cognitive networks were related to cognitive measures, both cross-sectionally and longitudinally. **A)** At baseline, higher energy of DorsAttn⟷Executive was related to better performance on the immediate memory subscale of the RBANS (top). We confirmed that the concurrent longitudinal changes in these brain and cognitive performance measures were related (bottom), so that an increase in energy cost of DorsAttn⟷Executive transitions accompanied a decline in immediate memory performance. **B)** Meanwhile, lower energy of Executive⟷Def+Lim was related to better performance on the attention subscale of the RBANS. There was also a concurrent longitudinal change in this transition cost and cognitive performance (bottom), so that a decrease in Executive⟷Def+Lim energetic cost was associated with a decline in attention performance. The concurrent change in brain and cognitive measures was estimated by modeling the individual slopes for time-related change within each participant separately for the cognitive and the biomarker variables and subsequently correlated. **C)** Higher level of Alzheimer’s disease pathology, pTau181, was related to facilitated transitions between antagonistic networks DorsAttn⟷Def+Lim at baseline. However, baseline level of pTau181 was not significantly associated with change in transition energy between these states (p=0.07). The uniform model is presented, but all other models showed the same relationship. Additionally, the models informed by the PET data also indicated a negative relationship between SomMot⟷Executive transition energy and pTau181 (p’s < 0.05).

Further, to determine whether the level of p-Tau181 biomarker measured at a single time point also predicted the *change* in the energetic cost of DorsAttn⟷Def+Lim transitions, we fitted an LMM with individual intercepts and individual slopes for the effect of time from baseline. The model was adjusted for covariates. However, this model was singular and was not further interpreted. Overall, these findings suggest that the relationship between AD pathology and brain energetics is not generalized across network reconfigurations, but may instead emerge selectively for transitions involving networks, such as the default network, that are particularly vulnerable to AD pathology^24^.

### Energetic costs of transitions between higher-order networks were related to cognitive performance

Similarly, we hypothesized higher energetic costs of transitions to be associated with greater costs of state transitions. More specifically, we expected these effects to be specific to higher-order associative, and transmodal networks engaged in complex cognition (executive, dorsal attention, ventral attention, default and limbic), rather than the sensory and motor unimodal networks.

Therefore, grounding the energetic costs of state transitions in the context of cognition performance, we fitted LMMs where each cognitive subscale of the RBANS as well as the total RBANS score were modeled individually. The energetic cost of each transition was used as the main predictor in separated models. Time from MRI baseline visit, the previously used covariates, and the individual intercepts were included. The tests were FDR-corrected for 16 comparisons within each cognitive domain.

Higher energetic cost of transitions between DorsAttn⟷Executive states was associated with better performance on the immediate memory subscale of the RBANS (Figure 4B). Meanwhile, lower cost of Executive⟷Def+Lim transitions was related to better performance on the RBANS’ attention subscale (Figure 4C). Subsequently, we assessed whether these relationships held longitudinally. Individual slopes for MRI and RBANS data were extracted and correlated.

Energetic costs were fitted using models predicted by time from baseline MRI, covariates, individual intercept, and individual slope for time from baseline MRI. RBANS scores were fitted using models predicted by time from baseline RBANS assessment, covariates, individual intercept, and individual slope for time from baseline RBANS assessment.

We confirmed that longitudinally immediate memory declined alongside increased cost of DorsAttn⟷Executive transitions (Figure 4B), and that attention declined concurrently with a decrease in Executive⟷Def+Lim energetic cost (Figure 4C). Together, these findings suggest that cognitive decline may be accompanied by selective alterations in the energetic requirements for transitions between the higher-order large-scale brain networks rather than reflecting a global loss of network efficiency.

### Sex differences in energetic costs of state transitions and their relationships to Alzheimer’s disease pathology and cognition

Females and males differ in the rate of decline of white matter with females showing accelerated decline in some white matter tracts compared to males^25^. The sexes also show differences in metabolic changes relevant to development of AD^26^. As these factors are thought to affect the energetic costs, sex differences were expected, with augmented effects in females compared to males. To assess the effects of sex assigned at birth on energetic costs in older adults, we compared the costs of state transitions between females and males at baseline. Linear models were fitted predicting the energy of each transition with sex assigned at birth and adjusting for covariates. Of note, the sample contained more assigned female (N=192) than assigned male (N=81) participants, with females comprising 70% of the sample. Females showed greater baseline costs of Visual⟷Def+Lim and Visual⟷Executive transitions, compared to males. Meanwhile, males had larger baseline energy costs of SomMot⟷Executive transitions, compared to females (Figure 5).

**Figure 5.**
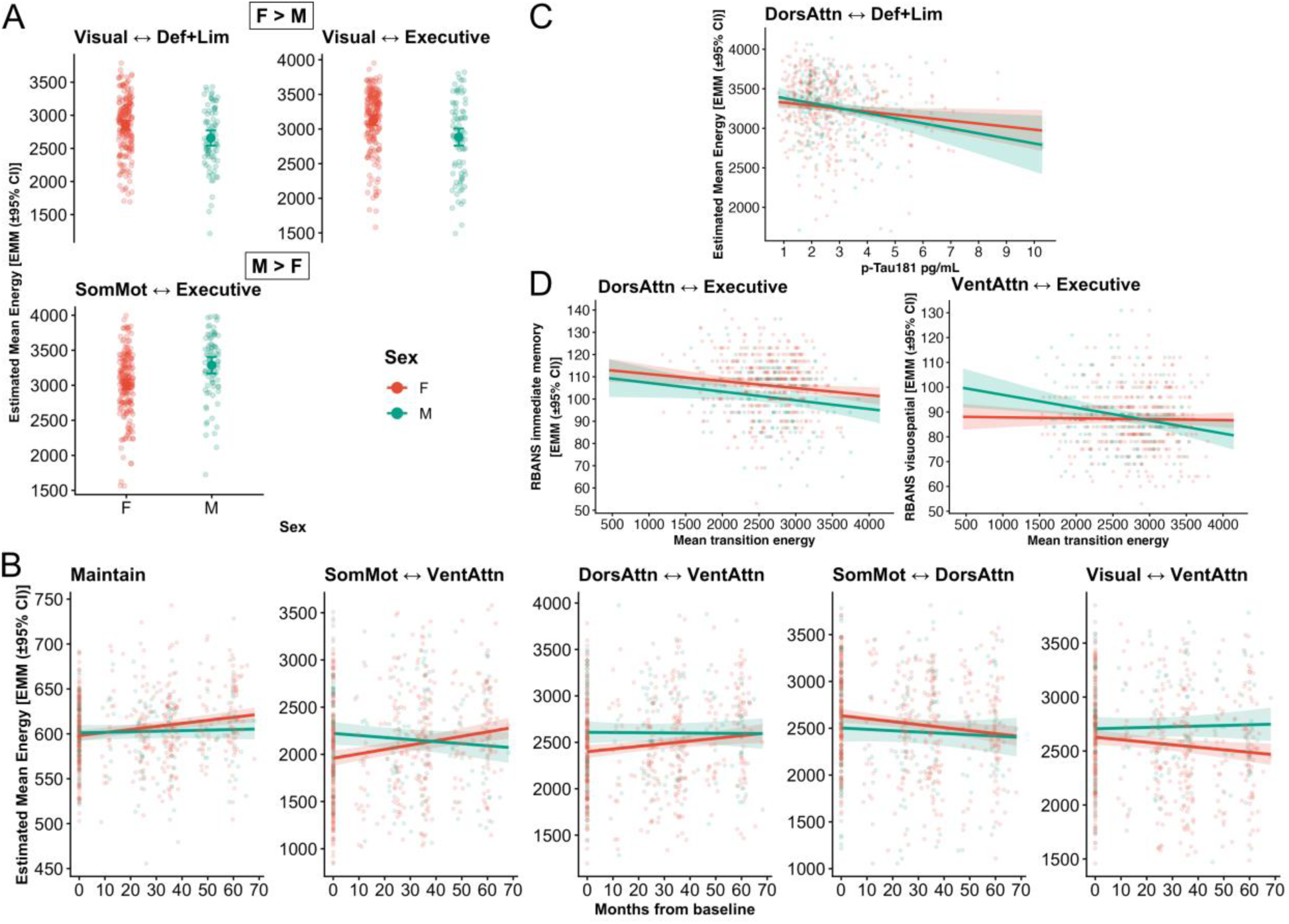
Assigned sex might affect the energy cost of transitions. We investigated the baseline differences in transition costs between females and males and repeated the main set of analyses on data disaggregated by assigned sex. **A)** There was a sex-related divergence in transitions associated with different sensory and motor domains. Females showed greater baseline costs of Visual⟷Def+Lim and Visual⟷Executive transitions, compared to males, while males had larger baseline energy costs of SomMot⟷Executive and transitions, relative to females. **B)** Within the subsample of females we observed increased energetic costs of maintenance, SomMot⟷VentAttn, and DorsAttn⟷VentAttn transition costs, and a decrease of energetic costs associated with SomMot⟷DorsAttn and Visual⟷VentAttn transitions. Meanwhile, none of the slopes were significant in the relatively small sample of males. **C)** The association between DorsAttn⟷Def+Lim transition cost and p-Tau181 was significant for males but not females. **D)** The association between DorsAttn⟷Executive transition cost and immediate memory performance was significant in females only. Unlike in the whole sample, the Executive⟷Def+Lim transition cost was not significantly related to cognition in the data disaggregated by assigned sex. Meanwhile, there was a significant relationship between low cost of VentAttn⟷Executive transitions and better performance on the visuospatial constructional scale of the RBANS in males but not in females. Note that the cohort consisted of an uneven number of assigned female (N=192) and assigned male (N=81) participants, with females comprising 70% of the sample. The observed trends that will require confirmation in larger, sex-balanced samples.

Previously described analyses were repeated on data disaggregated by assigned sex. The LMMs models were adjusted by removing sex from the list of covariates. Only the main uniform model was tested. First, we assessed whether there was a significant longitudinal change in the energetic cost of transitions for females and males separately. Within the subsample of females we observed increased energetic costs of state maintenance, SomMot⟷VentAttn, and DorsAttn⟷VentAttn transitions. Unlike in the entire sample, the cost of Visual⟷SomMot transitions was not increased among just females. Meanwhile, DorsAttn⟷VentAttn transition cost was not significantly increased in the total sample. We also observed a decrease of energetic costs associated with SomMot⟷DorsAttn and Visual⟷VentAttn transitions. The latter decrease was not significant in the entire study sample. The effect of time from baseline was not significant in the subgroup of males for any transition. The effects of time from baseline for the sex-disaggregated NCT models are reported in Table 2.

Next, we investigated the relationships between the energetic costs and AD risk and pathology. The negative association between DorsAttn⟷Def+Lim transition cost and p-Tau181 was significant for males only. No additional relationships were identified. The association between the energetic costs and cognitive performance on the RBANS was also investigated for each assigned sex separately. The cost of the DorsAttn⟷Executive transition was cost significantly related to immediate memory performance in females, but not in males. The Executive⟷Def+Lim transition showed a significant relationship to attention subscale in the complete dataset, but not in the in the data disaggregated by sex. The concurrent longitudinal associations were not confirmed. Overall, these findings suggest that energetic changes accompanying aging are not uniform across network transitions and may have distinct cognitive and pathological correlates across sexes, highlighting the potential importance of considering assigned sex when examining how altered network reconfiguration relates to cognitive aging and AD risk.

## Discussion

In this longitudinal study in a cohort of older adults at familial risk of Alzheimer’s disease, we leveraged multi-modal neuroimaging data to investigate aging-related changes in the energetic costs associated with maintaining network-based brain states and with transitions between these states using the computational Network Control Theory (NCT) model. In agreement with our hypothesis, we found increased energetic costs of state maintenance and some between-network transitions but also cost decreases. Specifically, aging affected the energetic costs of transitions between the sensorimotor and the attention states, with increased costs for the unimodal states (Visual⟷SomMot), between SomMot⟷VentAttn, and between the attention states (DorsAttn⟷VentAttn) but only in females. On the other hand, the cost of SomMot⟷DorsAttn transitions decreased in the whole sample, while the cost of Visual⟷VentAttn transition decreased in females only. Although the inclusion of cortical features in the models resulted in lower transition costs, they did not substantially affect the trajectories of change. We also explored relationships of these costs to both Alzheimer’s disease pathology and risk factors, and cognitive function. Contrary to our expectations, greater cost of transition was not necessarily related to more pathology or worse cognition. Instead, the relationship depended on the specific pair of networks involved. We saw higher levels of AD biomarker p-Tau181 predicted less costly transitions between DorsAtten⟷Def+Lim networks. Meanwhile, more costly (or increased cost of) transitions between Executive⟷DorsAtten networks were associated with poorer (or decreased) immediate memory performance, while easier (or decreased cost of) transitions between Executive⟷Def+Lim networks were linked to a lowering in attention scores. Together, our findings suggest that aging-related changes in brain energetics are not adequately characterized by a global increase in the difficulty of reconfiguring brain states. Rather, they reveal a selective reorganization of the energetic requirements for coordinating particular functional systems, with distinct transitions showing different relationships to AD pathology and cognitive function. By quantifying the energetic demands of moving between network states, our NCT approach provided a complementary perspective on brain aging that captures aspects of network reconfiguration not apparent from measures of connectivity or network activity alone.

### Brain dynamics of sensory, motor and attentional systems undergo transition-specific reorganization in aging

There were increases in control energy of state transitions with time, but decreases were also observed, suggesting that aging is related to an overall reorganization rather than a uniform increase in the energetic demands of brain dynamics. The increases may reflect greater resistance imposed by age-related degeneration of structural pathways, consistent with longitudinal diffusion MRI evidence of white matter deterioration with aging^27^. This interpretation was partly supported by the IS/GF model, in which individual differences in energetic costs arose from variability in the structural *A* matrices while functional states were group-averaged. Costs of all transitions increased with time in this model, but not in the main model, suggesting that structural aging may impose broad constraints on network control. Conversely, the GS/IF model produced a pattern similar, albeit attenuated, to the main model, indicating that individual functional configurations also substantially contribute to which transitions become energetically altered. State maintenance was an exception, with the main model more closely resembling IS/GF. Because maintenance involves the same initial and target state, its energetic cost may depend primarily on the structural constraints supporting persistence rather than on the specific functional pattern expressed by each participant. The increased cost of state maintenance is consistent with evidence of reduced neural stability and impaired persistence of well-organized brain states in older adults^28,29^. Together, these findings extend prior work on age-related changes in structural connectivity, network organization, and brain-state dynamics^2,12,30^ by demonstrating that the energetic demands of network reconfiguration change longitudinally and differ across specific state transitions.

Apart from state maintenance, the longitudinal effects involved transitions between unimodal sensory/motor and attentional networks. This pattern is consistent with evidence that aging alters the integration of sensory and higher-order systems, with reduced network segregation and increased between-network connectivity accompanying age-related changes in large-scale functional organization^2^. Dynamic approaches similarly indicate age-related alterations in brain-state transitions and reduced flexibility, with the ease of transitions between some higher-order networks associated with cognitive performance^1,30^. Notably, recent work combining functional dynamics with NCT found that the probability of the unidirectional transition from the Visual/SomMot state to VentAttn/DorsAttn state was higher in older than in younger adults with the node-wise pattern of control energy of that transition showing relevance to visual working memory and age^12^. However, as the sensory/motor and attentional states were combined, it is unclear if these results are contradictory to the current findings showing aging-related increase in the SomMot⟷VentAttn.

The increases in transition energies were not accompanied by changes in their corresponding probabilities, which might suggest the aging brain maintains the ability to effectuate the transitions despite the increased energetic costs. This may be particularly relevant for attentional systems, as aging has been associated with altered functional organization and reduced cognitive flexibility, including changes in the coordination and lateralization of attention-related networks^31^. The decrease in transition efficiency between the VentAtten and DorsAtten networks observed in females might imply a reconfiguration between distinct attentional modes, affecting the ability to flexibly switch between stimulus-driven (VentAttn) and goal-directed (DorsAttn) attentional processes. However, systematic cohort-level changes in energetic cost did not correspond to robust individual differences in RBANS performance, potentially reflecting the limited sensitivity of broad neuropsychological measures to subtle alterations in early sensory-attentional processing.

### Individual differences in energetic cost of transitions between higher-order cognitive networks relate to AD pathology and cognition

Cognitive trajectories of change, as well as AD pathology, were related to transitions between the higher-order cognitive networks: Executive, Def+Lim and DorsAtten. Higher levels of blood plasma pTau181 were related to facilitated transitions between antagonistic networks DorsAtten⟷Def+Lim, which typically show reduced anticorrelation in aging and dementia^32,33^. Easier transitions between these networks, which were most effortful compared to other transitions at baseline, may suggest pathological merging of these distinct states in those with high AD burden.

Surprisingly, we did not observe differences in energetic costs between the APOE groups, expected due to APOE4 allele’s role in brain energy metabolism^34,35^ and myelin pathology^22,36^. The APOE4 groups did not show differences in structural connectivity in our sample which might explain the lack of effect. Furthermore, glucose metabolism is also affected by maternal (but not paternal) family risk of AD^35^ present in 72% of the sample, possibly obscuring the effect of the APOE4 gene on energy metabolism. The energetic costs were also not sensitive to cognitive vulnerability defined by long-term decline on the RBANS^37^. However, cognitive vulnerability was defined using the entire RBANS battery, whereas the presented effects were domain- and transition-specific.

Immediate memory and attention domains showed coupled longitudinal change with energetic costs of transitions involving the executive network. Its facilitated transitions to/from the dorsal attnention network were associated with better memory performance. Although network transitions and coupling are not equivalent, efficient transitions between these networks might similarly support performance on task requiring focused attention^38^. Although Executive-Def+Lim coupling might support certain type of cognition^39^, especially in older adults^40,41^ increased communication between these systems might be detrimental to externally-oriented attention tasks^42^. This possibility was supported by our observation of facilitated Executive⟷Def+Lim transitions (which already required the least energy compared to other transitions at baseline) being associated with worse attention performance.

### Future directions for NCT models in aging and AD

Therefore, the presented results fit into and enrich the current understanding of neurocognitive aging, supporting the value of NCT modeling. However, energetic costs showed limited sensitivity to AD-related pathology and risk factors, apart from the one relationship with p-Tau181. The inclusion of PET-derived cortical pathology did not seem to affect the trajectories of transition costs. However, cross-sectionally, we observed a universal reduction of the energetic costs in these weighted models compared to the uniform model. This should not be interpreted as presence of pathology “facilitating” the transitions. Previous work suggests that inclusion of biologically meaningful weights, such as receptor distributions^9^ can reduce estimated energetic costs, possibly providing a more accurate representation of brain dynamics, especially if the feature follows major cortical gradients.

Future work might assess how regional pathology affects the node-level energetic inputs based on its placement in the organizational hierarchy. Previous NCT work has shone light onto energetics of transitions across cytoarchitectonic hierarchy in youth^19^. The study found that bottom-up transitions were easier to complete than the top-down ones, which might go against our observation that unimodal states were easier to transition into than out of, while the preferred direction was reversed for the transmodal states. The methodologies of the studies are not directly comparable as we did not separate the origin and the target for the inbound and outbound transitions. However, it is possible that there is a lifespan shift in energetic asymmetries, to be assessed in future studies.

Further investigation into the sex differences in control energies are also warranted. We have shown that the cost of DorsAttn↔VentAttn transitions increases in females but not men, possibly supported by a recent finding that females have lower functional connectivity between these two networks than males regardless of age^43^. Another study in youth found sex differences in controllability^44^, but it is unknown if they persist into older adulthood. Future research designed with sex differences in mind could further investigate whether females and males differ in energetic costs during aging. Given the sex differences in the decline of white matter^25^ and metabolic changes relevant to development of AD^26^, such differences are expected.

In conclusion, by leveraging multi-modal MRI and PET data we build a uniquely interpretable NCT model, which allowed for the first longitudinal characterization of energetic costs in older adults at familial risk of AD, with grounding in concurrent cognitive change and p-Tau181 pathology accumulation. NCT was able to capture the reorganization of brain dynamics predominantly affecting the transitions involving sensorimotor and attention networks, with hypothesized increases in energetic costs of transitions, but also an unexpected decrease in cost. Aside from cohort effects, we observed individual differences in concurrent changes in the energetic costs of transitions between the higher-order networks and cognition. The study contributes to the growing literature of network control and other dynamic models of AD^45–47^, providing novel insight into the developing energetic landscape of the aging adult brain.

## Methods

### Participants

This study used a subset of participants from the PREVENT-AD (Pre-symptomatic Evaluation of Experimental or Novel Treatments for Alzheimer’s Disease) cohort^48,49^. The PREVENT-AD project, which has been collecting data since 2011, provides access to some of its acquired data in an open repository (https://openpreventad.loris.ca). Participants in the PREVENT-AD study were older adults with a first-degree familial history of Alzheimer’s disease, making them at elevated risk of developing the disease. 72% of participants had maternal family history of AD, 38% had paternal family history of AD and 13% had had both parents affected by the disease. In 4% only the sibling or siblings were affected. All participants were cognitively healthy at recruitment. However, some of them developed MCI during the course of the study. For the presented analyses, data of 279 adults was included, with 128 participants completing all three MRI sessions. Seven participants were missing at least one demographic variable. Therefore, the statistical models where demographic information were used as covariates included data of 272 participants with 603 observations in total. From the participants with available ethnicity data 269 were white and one was black, while three participants identified as Hispanic. Ethnicity was not further used as a covariate in any models. The sample’s ethnic distribution is consistent with the majority of Montreal’s older adult population. Over half of the sample were females, which is consistent with greater longevity of females compared to males. For the presented analyses, data of 279 adults was included, with 128 participants completing all three MRI sessions. PET data was available for 188 participants at a single time point. PET scans were not performed on the same day as the MRI scans. See Table 1 for demographic and cognitive details. Before participating in the study, all participants gave informed written consent. The procedures of the PREVENT-AD study were approved by the McGill Institutional Review Board and/or the Comité d’éthique de la recherche du CIUSSS de l’ouest de l’île de Montréal.

### Neuroimaging Acquisition

#### MRI Protocol

Multi-modal magnetic resonance imaging (MRI) data was acquired on a 3 T Siemens Magnetom PrismaFit MRI scanner equipped with a 32-channel head coil. The scanning protocol included structural scans: 1) high-resolution anatomical T1-weighted MPRAGE (magnetization-prepared rapid acquisition gradient echo) and 2) MP2RAGE^50^, 3) multi-shell 2D diffusion weighted imaging (DWI) acquired using spin echo echo-planar imaging (EPI), 4) multi-parametric mapping (MPM)^51^ using a gradient echo (GRE) sequence, 5) B1+ transmit field maps, as well as 6) a multi-echo functional MRI scan with a multiband gradient-echo EPI sequence for blood oxygenation level-dependent (BOLD) signal. The custom MPM sequence was developed and provided by the McConnell Brain Imaging Centre at the Montreal Neurological Institute.

1) **MPRAGE**: 1-mm isotropic resolution, TR = 2300 ms, TE = 2.96 ms, TI = 900 ms, TA = 5:30 min (repetition time, echo time, inversion time, acquisition time, respectively), flip angle = 9°. This image was used segmentation of the brain tissue, extraction of the cortical surface, and registration to the atlas.
2) **MP2RAGE**: 1-mm isotropic resolution, TR = 5000 ms, TE = 2.76 ms, TI1 = 940 ms, TI2 = 2830 ms, TA = 7:07 min, flip angles = 4°, 5°. The image was used to derive T1 and equilibrium magnetization (S0) for computation of MTsat maps. S0 maps were scaled by a gain factor of 2.5 to match the receiver gain of the MT-weighted acquisition prior to MTsat estimation.
3) **DWI**: 2-mm isotropic resolution, TR = 3000 ms, TE = 66 ms, posterior-anterior (PA) phase encoding, TA = 5:49. Across three shells, 109 measurements were taken isotopically spaced around a sphere with the following *b*-values: 9 × *b* = 0, 7 × *b* = 300, 29 × *b* = 1000, 64 × *b* = 2000 s/mm^2^. Additionally, 5 × *b* = 0 images were taken with reversed (anterior-posterior; AP) phase encoding direction and identical sequence parameters for the purpose of distortion correction. The DWI images were used for whole-brain tractography and mapping of the intra-cellular volume fractions.
4) **MPM** : 1-mm isotropic resolution, TR = 27ms, six TEs = 2.04-14.89 ms, echo-spacing = 2.57 ms, flip angle = 6°. This included an MT-weighted, proton density-weighted, and T1-weigthed image for the purposes of computing MTsat. The MT pulse had the following parameters: TR = 27 ms, six echoes, TEs = 2.04–14.89 ms, echo-spacing = 2.57 ms, flip angle = 6°.
5) **B1+:** two spin-echo echo-planar sequences with the same 2 × 2 × 4 mm resolution, TR = 4010 ms, TE = 46 ms, TA = 1:08 min, and different with flip angles = 60°, 120° to compute a B1+ map.
6) **fMRI-BOLD**: Whole-brain functional images were collected in the axial (transversal) plane with 48 interleaved slices (slice thickness = 3.0 mm; no slice gap) covering the entire brain. Imaging parameters were as follows: TR = 1000 ms; three TEs = 12.00, 30.11, and 48.22 ms; TA = 10:24 mins, flip angle = 50°; field of view (FOV) = 240 × 240 mm²; acquisition matrix = 80 × 80 (in-plane resolution = 3.0 × 3.0 mm²); anterior-posterior (AP) phase encoding phase encoding direction. Simultaneous multi-slice (multiband) acceleration with a factor of 4 and in-plane GRAPPA parallel imaging with an acceleration factor of 2 (24 reference lines) were used to reduce acquisition time and susceptibility-related distortions. A total of 604 volumes were acquired.

### PET acquisition and processing

Amyloid (Aβ; [18F]NAV4694) and tau ([18F]flortaucipir) PET scans were acquired. Amyloid-PET images were taken 40-70 minutes after injecting ∼6 mCi of the radiotracer, while the tau-PET images were acquired 80-100 minutes after injecting a dose of ∼10 mCi, in five-minute frames. PET scanning was performed at the McConnell Brain Imaging Centre at the Montreal Neurological Institute using a brain-dedicated Siemens/CTI high-resolution research tomograph scanner. We used single time-point data from the first wave of data acquisition that took place between 2016-2017.

The preprocessing was previously described in^52^, with the pipeline published here: https://github.com/hayleyshanks/Longitudinal-MRI-PET-preproc. However, instead of the Desikan-Killany atlas, the Schaefer atlas was used to obtain the volume-weighted mean of the standardized uptake value ratio (SUVR).

### MR data preprocessing

#### Structural data

The structural MRI preprocessing pipeline *micapipe* (v0.1.5)^53^ was adapted to preprocess the anatomical and the diffusion data to obtain the structural connectomes. The full preprocessing pipeline is described in detail in^54^. In brief, Structural T1-weighted images were processed using FreeSurfer (v6.0)^55^ for cortical reconstruction and segmentation, while subcortical structures and tissue segmentation were obtained using FSL FIRST and FAST. The resulting five-tissue-type segmentation was used for anatomically constrained tractography. A previously validated segmentation procedure of the T1w images was used to quantify white matter hypointensities^56,57^. A single extreme outlier was excluded from the sample (z-statistic = 13.4).

Diffusion-weighted images were preprocessed in MRtrix3 (v3.0.3)^58^, including denoising^59^, Gibbs ringing removal^60^, correction for susceptibility-induced distortions, head motion and eddy currents using FSL TOPUP/eddy^61–63^, and B1 bias-field correction^64^. The diffusion data were upsampled to 1-mm isotropic resolution, followed by multi-shell, multi-tissue constrained spherical deconvolution and intensity normalization. Anatomically constrained probabilistic tractography (iFOD2) was then performed to generate whole-brain tractograms containing 1 million streamlines.

Semi-quantitative myelin-sensitive MTsat maps were calculated using the hMRI toolbox (v0.5.0)^65^ from the MT-weighted MPM acquisition, including B1+ correction using the double angle method. The procedure is further described in^66^. The MTsat maps and diffusion-derived neurite density maps were combined to estimate myelin volume fraction (MVF) and intra-axonal volume fraction (AVF) using a calibrated aggregate g-ratio framework^54^. Streamlines were subsequently filtered using COMMIT^67^ to estimate tract-specific myelin and axonal cross-sectional areas. Finally, structural connectivity matrices weighted for total intra-axonal cross-sectional area were generated in the Schaefer-400^68^ cortical parcellation. No further filtering of the resulting structural connectomes was performed.

#### Functional data

Multi-echo resting-state fMRI data were preprocessed using the ME-ICA pipeline (v3.9.3), which combines AFNI- and TEDANA-based processing to separate BOLD from non-BOLD signal components^69,70^. Preprocessing included despiking, slice-timing correction, deobliquing, axial reorientation, rigid-body motion correction (using the first echo as the reference), and co-registration to the participant’s T1-weighted anatomical image, followed by nonlinear normalization to MNI space. Motion correction parameters estimated from the first echo were applied to all echoes, and a brain mask was generated from voxel-wise T2* and S0 maps estimated from the multi-echo data. Functional images were intensity normalized prior to denoising. Spatial smoothing was not applied. The three echoes (TE = 12.0, 30.11, and 48.22 ms) were optimally combined using voxel-wise T2* weighting, after which TE-dependent independent component analysis (ME-ICA) was performed with TEDANA to identify and remove non-BOLD components while retaining BOLD-related signal. The BOLD components were used to reconstruct the optimally combined, denoised time series with the thermal noise removed (high κ [kappa] data). These resulting time series in native anatomical space were parcellated into 400 cortical region time series using the Schaefer atlas^68^ and used for subsequent analyses.

### Creation of brain states

We defined network-based “brain states” according to the approach described in^20^ based on their correspondence to Yeo-Krienen^71^ intrinsic resting-state networks. Briefly, to obtain the states, the preprocessed fMRI time series from the 400 Schaefer cortical regions were z-scored across time, and the average time series was calculated for each network, resulting in six time series. At each time point, the network with the highest average activation was deemed dominant and assigned as the state for that time point. A fixed threshold (t = 10) was used to discard time points that did not reach sufficient activation. We then averaged the time points dominated by the same network to obtain the state maps, which were subsequently input into the NCT models.

The networks were: Visual, Somatomotor (SomMot), Dorsal Attention (DorsAttn), Ventral Attention (VentAttn), and Executive (also known as Control or Fronto-Parietal network), and Default and Limbic (Def+Lim) combined into a single network. The Default and Limbic states were combined a priori based on evidence that regions traditionally classified as the Limbic network, particularly orbitofrontal and anterior temporal cortices, form part of an extended Default network^72^. The separation of these regions as a distinct Limbic network has been attributed in part to susceptibility-related signal loss in single-echo acquisitions affecting orbitofrontal and anterior temporal regions^72^. Multi-echo fMRI, such as the one used in this study, can improve BOLD sensitivity in regions affected by such signal loss; accordingly, we combined the Default and Limbic regions into a single network for the present analyses.

### Network Control Theory

#### Model formulation

Network control theory (NCT) was used to estimate the control energy required to transition between macroscopic functional brain states, given constraints imposed by the brain’s structural connectivity^73,74^. The structural network was represented by an adjacency matrix *A,* in which nodes corresponded to brain regions and edges represented white matter bundles.

In conventional applications of NCT to DWI data, edge weights are derived from tractography measures such as the number of streamlines (NOS). In the current study, we instead incorporated microstructural information afforded by MTsat maps to estimate tract caliber. Accordingly, each edge in *A* represented the total intra-axonal cross-sectional area of each tract, increasing the biological plausibility of the model. Before modeling brain state dynamics, A was normalized to enact a particular form of dynamical stability: 1) 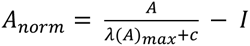, where *λ*(*A*)*max* is the largest eigenvalue of *A* and *c* is a constant, here set to 1, that determines if the system approaches the corresponding mode of *A* or decays.

Brain states were modeled as a linear time-invariant system, following: 2) ***ẋ*** = *A* × ***x***(*t*), where ***x***(*t*) is the vector of activity states across *N* brain regions. Thus, the instantaneous evolution of the brain state is dependent upon its current state and the time-invariant anatomical connections encoded by *A*. A linear model was adopted on the basis of previous work demonstrating that linear systems adequately model macroscale brain dynamics while remaining computationally tractable. Of note, noise is not explicitly modeled in this approach^74,75^.

Equation 1 presents the system evolving in the absence of external control. To model transitions between brain states, an external control input **u**(*t*) was introduced, such that the equation is extended to 2) **ẋ** = *A* × **x**(*t*) + *B* × **u**(*t*). Here, *B* is a matrix whose non-zero diagonal elements indicate which brain regions are receiving control input. An identity matrix *B* provides a full and uniform control set, whereas setting a diagonal element to zero would reduce its control input to zero. Diagonal values of *B* above 0 and below 1 provide weights to the control inputs. *A* and *B* are N × N matrices while **x** and **u** are N × 1 vectors, where N stands for the number of controllable brain regions or voxels.

For each transition, the system was required to evolve from an initial brain state **x**(0) to a target state **x**(*T*) over a specified time horizon *T.* The optimal control input was identified by minimizing a cost function that penalized both deviation of the state trajectory **x** from the target state and the magnitude of the control input: 3) 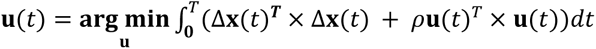, where Δ**x**(*t*) = **x**(*T*) − **x**(*t*) is the distance between the target state **x**(*T*) and **x** at any given time *t*, and *ρ* (rho, also referred to as the mixing parameter) determines the relative cost assigned to the control signals and the state trajectory. It is typically set to 1, as was done here, thereby assigning equal cost weighting to these two components.

Following the minimization of **u**(*t*), the optimal control energy *E* required to complete the transition was calculated by integrating the squared magnitude of the control input over the time interval: 4) 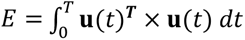.

In depth discussion of the derivation of the optimal control framework and the effects of alternative model parameters can be found in Ref ^74^.

#### Estimation of control energy for state transitions

Here, we computed the energy costs for transition trajectories between all pairs of six states obtained from the functional MRI data (see *2.3.3. Functional brain states*), including symmetric transition pairs (e.g., Visual⟷SomMot and SomMot⟷Visual) and self-transitions (e.g., Visual⟷Visual). Such symmetric transition pairs were averaged given very high correlation between them (r≈.99). We also averaged all self-transition costs into one *maintenance cost* (sometimes referred to as “persistence” energy). This resulted in 16 average costs in total. Next, we obtained the difference, or asymmetry, between all the average inbound (e.g., *any network except Visual* → Visual) and all the average outbound (e.g., Visual → *any network except Visual*) transitions for each state. Initial and target state vectors were normalized before optimization.

In total we tested six models; in four of these models, individual structural connectomes and individual functional states were used to obtain the energy costs for each participant. Among these individual-data models, we varied the control scheme, such that the *B* matrix either encoded uniform control (equal control weights of 1 for all nodes), control weighed by cortical features (cortical thickness derived from T1w structural data after Freesurfer processing), amyloid-PET, or tau-PET (see *PET acquisition and processing*). This allowed for testing the effect of incorporating the spatial distribution of these cortical features on the age-related trajectories of control energy. Further, to control for the influence of structure vs functional state on control energy estimates, we also tested models with uniform control where either the structural connectome or the functional states were group averaged before estimation of the model, so that energy costs of each participant did not use individualized structural or functional data. The subject-level transition energies from each model were subsequently used for statistical analyses. All models were estimated using the *nctpy* toolbox^76^ for Python.

### Calculation of transition probability and its relationship to energetic costs

Transition probabilities were calculated from the subject-specific sequence of functional brain states identified at each fMRI time point (TR) (see *Creation of brain states*). For each participant, every pair of consecutive time points was treated as a state transition, including transitions in which the brain remained in the same state (self-transitions/state maintenance). The number of observed transitions from state *i* at time *t* to state *j* at time *t* + 1 was counted for all possible state pairs, yielding an asymmetric 6 × 6 transition count matrix (after ignoring the states that did not meet the specified activation threshold). Counts within each originating state were then normalized by the total number of transitions originating from that state to obtain conditional transition probabilities. Note that the rows and column do not sum to 1 due to the removal of the non-state time-points. The group-averaged transition probability matrix is shown in Figure S1A. We used repeated measures correlations (*rmcorr* R package) to relate transition costs to transition probabilities, while accounting for multiple time points per participant. The symmetric transitions and self-transitions were not averaged for this analysis to respect the directionality of the transitions, resulting in 36 tests.

### Behavioral data: Cognitive battery

The Repeatable Battery for the Assessment of Neuropsychological Status (RBANS)^77^ was used to test cognitive performance at baseline and follow-up visits. RBANS is composed of five subscales, each comprised of 2-4 subtests, designed to assess cognitive domains of: immediate memory (List Learning, Story Memory), delayed memory (List Recall, List Recognition, Story Recall, Figure Recall), attention (Digit Span, Coding), language (Picture Naming, Semantic Fluency), visuospatial/ constructional ability (Figure Copy, Line Orientation). Age-adjusted index scores were used.

### Biomarkers of Alzheimer’s disease

We also assessed the relationships between the control energies and risk factors for AD, including APOE4 carriership status, cognitive vulnerability, and blood plasma tau levels p-tau181. Blood-based plasma p-tau 181 was analyzed and quantified using in-house single molecule array assays. See^49^ for details. The participants were genotyped for *APOE* using the procedure described in^48^. Participants carrying at least one APOE4 allele were deemed APOE4 positive, while those without an APOE4 allele were considered APOE4 negative. Cognitive vulnerability was defined by the long-term trajectory of decline on the RBANS using the entire available PREVENT-AD data set collected in years 2012-2024, including the time preceding neuroimaging data acquisition included in this study. These calculations were first described and used in ^37^.

### Statistical analyses

#### Baseline analyses

We compared the energetic costs associated with each state transition using linear mixed-effects models (LMMs) with individual intercepts, where the energetic cost was predicted by the transition between a pair of states or by state maintenance. The used formula was: *Mean energy ∼ transition + (1 | ID).* The non-weighted uniform model and the models weighted for the cortical features: cortical volume, amyloid-PET and tau-PET, were compared using the formula: *Mean energy ∼ control scheme + (1 | ID).* To calculate the transition asymmetry for each network, we averaged the costs of transitions from any other state into a given state (inbound energy cost) and the average cost of transitions from that state to any other state (outbound energy cost). Then, we calculated the difference, Inbound – Outbound, between these two costs to obtain the transition asymmetry. The difference between the asymmetries was modeled as: *Mean energy asymmetry ∼ network state + (1 | ID).* We also performed t-test’s to determine if the asymmetry is significantly different from zero. Baseline data was also used to determine the sex differences in transition energies. This was estimated using a linear model (*lm* function in R; not mixed-effects) separately for each transition with the following formula: *Mean energy* ∼ sex + *education years + baseline age + FD + eITV*. The covariates were mean centered. The statistical analyses were performed in R. The LMMs were estimated using the *lmer* and *lmerTest* packages, while the post-hoc comparisons were performed using the *emmeans* package.

#### Longitudinal analyses

Longitudinal associations between time since baseline MRI and transition energy were assessed separately for each network transition using LMMs with participant-level random intercepts to account for repeated observations. Models included the primary predictor of interest (months since baseline MRI) and covariates. To improve robustness to influential observations, inference was based on robust linear mixed-effects models (*rlmer*) estimated using M-estimation. Fixed-effect estimates and t-statistics were extracted from the robust models, while degrees of freedom were obtained from the corresponding standard linear mixed-effects models (*lmer*). Satterthwaite’s degrees of freedom approximation method was used to obtain the degrees of freedom using the *lmerTest* package. Note that the Satterthwaite degrees of freedom account for the estimated variance-covariance structure of the fitted mixed-effects model, which can result in different denominator degrees of freedom for models fitted to the same participant data. Two-sided p-values were calculated from the t-statistics using the estimated degrees of freedom, and statistical significance was assessed using thresholds of p < 0.05, with correction for multiple comparisons performed using the false discovery rate (FDR) procedure across network transitions. The correction was applied across the set of network transitions tested within each model. The covariates were: demographic variables (age at baseline, sex, education years), estimated total intracranial volume (eITV) and in-scanner motion measured by framewise displacement (FD): *Mean energy ∼ months since baseline MRI + sex + education years + baseline age + FD + eITV + (1 | ID).* The models disaggregated by sex did not include sex as a covariate.

#### Analyses of relationships between energetic costs, AD risk factors and pathology, and cognition

Additionally, to estimate the effect of the AD risk factors and pathology on the transition energies, they were tested in separate models in interaction with the time predictor: *Mean energy ∼ AD risk factor or pathology * months since baseline MRI * + sex + education years + baseline age + FD + eITV + (1 | ID).* We also estimated the relationships between the change in energy costs and concurrent change in cognitive performance on the RBANS subtests. For this purpose, two models were estimated, one for the energy cost measure, one for the cognitive measure, followed by the extraction of the individual slopes for time since baseline (differing for cognitive assessments and MRIs often happening on different days), that were then correlated: *Model for cognition X: RBANS score ∼ months since baseline cognitive assessment + sex + education years + baseline age + FD + eITV + (1 + months since baseline cognitive assessment | ID)*. Model for energy cost Y: *Mean energy ∼ months since baseline MRI + sex + education years + baseline age + FD + eITV + (1 + months since baseline MRI | ID)*. The outcomes and predictors were centered prior to model estimation. Models that failed to converge or were singular were ignored. These analyses were repeated on data disaggregated by sex.

## Acknowledgements

We thank the PREVENT-AD research team and study participants for their time and dedication in collecting the data used in this study. A full list of PREVENT-AD authors and collaborators can be found at https://preventad.loris.ca/acknowledgements/acknowledgements.php?date=[2025-09-01]. We also want to thank the Laboratory of Brain and Cognition at the Montreal Neurological Institute for feedback on the project. Special thanks to Anaelle Wicquart for her help with quality assessments. This project was made possible by financial support from the Alzheimer’s Association (AARG22-927100) and the National Institutes of Health (NIA R01 AG068563). R.N.S. is a Research Scholar supported by the Fonds de la Recherche du Quebec – Santé (FRQS).

## Author contributions

P.S.: Conceptualization, methodology, software, formal analysis, data curation, writing—original draft, writing—review & editing, visualization, A.W.: Methodology, software, data curation, writing—review & editing. D.O’S.: Methodology, software, data curation, writing—review & editing. I.S.: Methodology, writing—review & editing. K.B.: Methodology, writing—review & editing. I.R.L.: Methodology, investigation, writing—review & editing. C.L.T.: Methodology, supervision, investigation, resources, funding acquisition, writing—review & editing. G.R.T.: writing— review & editing. D.S.B.: Conceptualization, methodology, supervision, writing—review & editing. R.N.S.: Conceptualization, investigation, resources, writing— review & editing, supervision, project administration, funding acquisition.

## Competing interests

The authors declare no competing interests.

## Data availability

This study utilized data from the PResymptomatic EValuation of Experimental or Novel Treatments for Alzheimer’s Disease (PREVENT-AD) study^48^. Data availability for PREVENT-AD is governed by the Open Access protocols. For more information about PREVENT-AD, please refer to https://douglas.research.mcgill.ca/prevent-alzheimerprogram/.

## Code availability

This study utilized several open-access resources. The adapted *micapipe*^53^ pipeline (v0.1.5; https://micapipe.readthedocs.io/) was used in preparation of the structural connectomes. Diffusion images were preprocessed using the MRtrix3 toolbox^58^ (v3.0.3; https://www.mrtrix.org/). Multiparametric maps were processed using hMRI toolbox^65^ (v0.5.0) (https://hmri-group.github.io/hMRI-toolbox/) in MATLAB. The NCT models were estimated and partially visualized using the *nctpy* toolbox^76^ (v1.0.1; https://nctpy.readthedocs.io/) for Python. The datasets were prepared using the functions from the *tidyverse* (v2.0.0) package in R (https://github.com/tidyverse). The statistical models used the *lme4* (v1.1-37), *lmerTest* (v3.1-3), *robustlmm* (v3.3-3), and *emmeans* (v1.11.1), *rmcorr* (v0.7.0) packages for R. The graphs were created using the *ggplot2* (v4.0.3) package in R.

## Notes

### Competing Interest Statement

The authors have declared no competing interest.

