## Supplementary materials for "Longitudinal changes in control energy of brain networks in older adults at familial risk for Alzheimer’s disease"

### **Relationship between probability of transition and its energetic cost**

To gain a better understanding of what the energetic cost represents we correlated the energy cost of each transition with transition probability of between the corresponding networks. All correlations between for the between-state transitions were negative and significant after FDR correction (all p’s $\leq$ 0.001), with the r-values ranging from -0.45 to -0.20 [95% CIs, -0.53, -0.10]. Therefore, the probability of transitioning between two networks was inversely related to the cost of that transition. The relationship between the cost of maintaining a state and the probability of its maintenance varied between states. Maintenance of Visual (r=-0.23, p < 0.001) and SomMot (r=-0.22, p < 0.001) states showed the same negative and significant relationship as the between-states transitions, so that the probability of maintaining these states was inversely related to its maintenance cost. This relationship was not significant for VentAttn (r=0.06, p = 0.31), Executive (r=0.05, p = 0.32) and Def+Lim (r=0.07, p = 0.22) states. However, DorsAttn state showed a significant positive relationship between maintenance probability and its cost (r=0.16, p = 0.003), so greater probability of maintaining that state was associated with higher cost (Supplementary Figure 1).

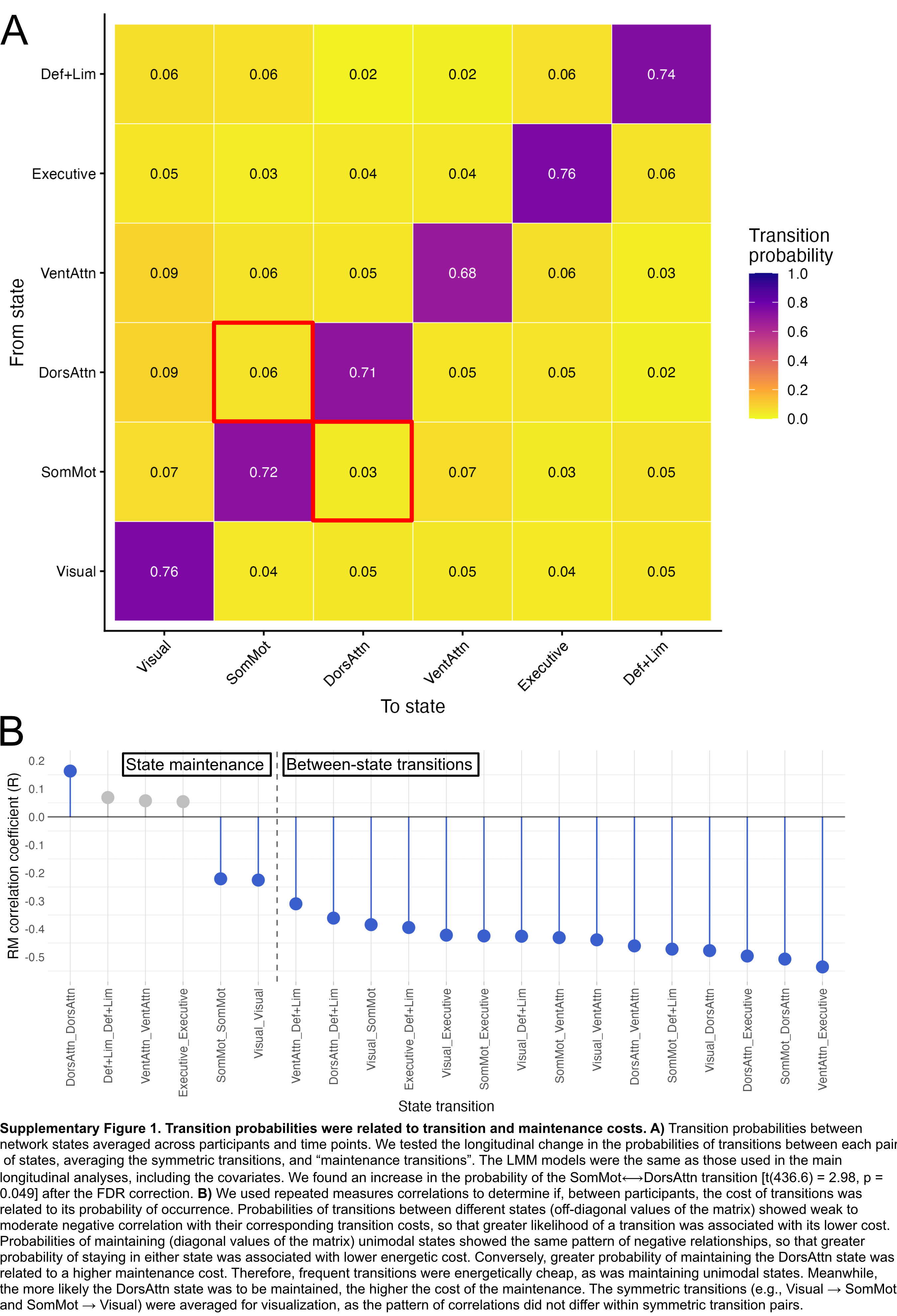

**Supplementary Figure 1. Transition probabilities were related to transition and maintenance costs. A)** Transition probabilities between network states averaged across participants and time points. We tested the longitudinal change in the probabilities of transitions between each pair of states, averaging the symmetric transitions, and “maintenance transitions”. The LMM models were the same as those used in the main longitudinal analyses, including the covariates. We found an increase in the probability of the SomMot⟷DorsAttn transition [t(436.6) = 2.98, p = 0.049] after the FDR correction. **B)** We used repeated measures correlations to determine if, between participants, the cost of transitions was related to its probability of occurrence. Probabilities of transitions between different states (off-diagonal values of the matrix) showed weak to moderate negative correlation with their corresponding transition costs, so that greater likelihood of a transition was associated with its lower cost. Probabilities of maintaining (diagonal values of the matrix) unimodal states showed the same pattern of negative relationships, so that greater probability of staying in either state was associated with lower energetic cost. Conversely, greater probability of maintaining the DorsAttn state was related to a higher maintenance cost. Therefore, frequent transitions were energetically cheap, as was maintaining unimodal states. Meanwhile, the more likely the DorsAttn state was to be maintained, the higher the cost of the maintenance. The symmetric transitions (e.g., Visual → SomMot and SomMot → Visual) were averaged for visualization, as the pattern of correlations did not differ within symmetric transition pairs.

### **Longitudinal changes in cognition**

Linear mixed-effects models were used to determine whether there was significant change in the RBANS, separately for the total score and each subscale. The main predictor was time from cognitive assessment baseline. The individual intercept was included alongside covariates: sex, baseline age, years of education and estimated total intracranial volume. The combined *rlmer* and *lmer* procedure was applied to obtain robust estimates, degrees of freedom and p-values, as described in the main Methods section (*Longitudinal analyses*). There was a significant longitudinal decline in total RBANS (t(403) = -5.09, p = < .001). Investigating the individual subscales, the decline was significant for: delayed memory (t(407) = -5.3, p = < .001), visuospatial constructional (t(445) = -5.87, p = < .001), and language (t(506) = -2.42, p = 0.0157), but not immediate memory (t(420) = -1.81, p = 0.0716) or attention (t(392) = 0.149, p = 0.881). Correction for multiple comparisons was not applied. Of note, the two RBANS subscales that showed significant individual difference in relation to energetic costs did not show age-related decline in the whole sample.

Supplementary Table 1: AD pathology and risk factor effects on transition energies

| network | p-Tau181 | APOE4+ | Cognitive vulnerability |
| --- | --- | --- | --- |
| Visual ↔ SomMot | 6.42, 0.905 | -4.62, 0.933 | 36.01, 0.840 |
| Visual ↔ DorsAttn | 27.65, 0.235 | 51.89, 0.571 | 95.23, 0.495 |
| Visual ↔ VentAttn | 15.27, 0.653 | 59.82, 0.571 | 60.82, 0.758 |
| Visual ↔ Executive | -12.93, 0.661 | -72.35, 0.517 | -39.65, 0.840 |
| Visual ↔ Def+Lim | -2.61, 0.905 | -62.86, 0.517 | -135.24, 0.230 |
| SomMot ↔ DorsAttn | -22.56, 0.534 | 85.95, 0.517 | 17.69, 0.962 |
| SomMot ↔ VentAttn | 5.43, 0.905 | -43.37, 0.650 | -96.66, 0.679 |
| SomMot ↔ Executive | -33.49, 0.212 | 33.98, 0.650 | -72.87, 0.679 |
| SomMot ↔ Def+Lim | 8.00, 0.905 | -11.18, 0.896 | 59.22, 0.758 |
| DorsAttn ↔ VentAttn | 37.32, 0.209 | 64.81, 0.517 | 139.32, 0.230 |
| DorsAttn ↔ Executive | -26.43, 0.235 | -64.55, 0.517 | -7.53, 0.962 |
| DorsAttn ↔ Def+Lim | -46.31, 0.011* | -11.68, 0.873 | -28.96, 0.840 |
| VentAttn ↔ Executive | 20.93, 0.534 | 35.58, 0.650 | 63.72, 0.758 |
| VentAttn ↔ Def+Lim | -24.96, 0.235 | -40.38, 0.622 | 5.15, 0.962 |
| Executive ↔ Def+Lim | 2.08, 0.905 | 30.99, 0.650 | 3.12, 0.962 |
| Maintain | -1.36, 0.653 | 4.09, 0.633 | 10.74, 0.380 |
| Values are β estimates and FDR-adjusted p-values. * pFDR < 0.05; ** pFDR < 0.01; *** pFDR < 0.001. Positive β’s indicate greater energetic costs are related to higher p-Tau181 levels, or greater energetic cost in APOE4 positive vs APOE4 negative participants, or greater energetic cost for participants showing cognitive vulnerability vs cognitively resilient participants. P-values were corrected using FDR. The energetic costs were extracted from the uniform model. | | | |

Supplementary Table 2: The effects on energetic costs on cognitive performance on the RBANS (Standardized β estimates and FDR-adjusted p-values)

| transition | Total | Immediate memory | Delayed memory | Attention | Visuospatial constructional | Language |
| --- | --- | --- | --- | --- | --- | --- |
| Visual ↔ SomMot | -0.24, 0.817 | -0.92, 0.586 | -0.56, 0.456 | 0.31, 0.900 | 0.33, 0.704 | -0.05, 0.984 |
| Visual ↔ DorsAttn | -0.29, 0.809 | 0.22, 0.969 | -0.15, 0.961 | 0.06, 0.937 | 0.04, 0.944 | -0.92, 0.442 |
| Visual ↔ VentAttn | 0.11, 0.903 | -0.69, 0.816 | -0.66, 0.456 | 0.25, 0.937 | 0.52, 0.652 | -0.02, 0.984 |
| Visual ↔ Executive | -0.54, 0.663 | 1.16, 0.547 | 0.09, 0.961 | -1.45, 0.307 | -1.56, 0.282 | 0.01, 0.984 |
| Visual ↔ Def+Lim | -0.24, 0.817 | -0.38, 0.958 | 0.79, 0.456 | -0.74, 0.673 | 0.16, 0.862 | 0.06, 0.984 |
| SomMot ↔ DorsAttn | 0.69, 0.663 | -0.34, 0.958 | 0.12, 0.961 | 0.96, 0.406 | 0.63, 0.592 | 0.86, 0.442 |
| SomMot ↔ VentAttn | 0.53, 0.663 | 0.07, 0.969 | 0.77, 0.456 | 0.11, 0.937 | 1.05, 0.377 | 0.49, 0.984 |
| SomMot ↔ Executive | 0.63, 0.663 | -0.39, 0.958 | -0.06, 0.961 | 1.31, 0.395 | 0.99, 0.472 | 0.37, 0.984 |
| SomMot ↔ Def+Lim | 0.40, 0.757 | 0.68, 0.816 | -0.36, 0.790 | 0.06, 0.937 | 0.34, 0.704 | -0.13, 0.984 |
| DorsAttn ↔ VentAttn | -0.06, 0.903 | -0.60, 0.816 | -0.23, 0.916 | 0.86, 0.465 | 0.65, 0.592 | -0.36, 0.984 |
| DorsAttn ↔ Executive | -1.08, 0.663 | -2.32, 0.018* | -0.66, 0.456 | -0.99, 0.465 | 0.36, 0.704 | -0.20, 0.984 |
| DorsAttn ↔ Def+Lim | -0.10, 0.903 | 0.19, 0.969 | -0.45, 0.786 | -0.74, 0.700 | -0.58, 0.652 | 0.57, 0.984 |
| VentAttn ↔ Executive | -0.61, 0.663 | 0.08, 0.969 | -0.63, 0.456 | -0.18, 0.937 | -1.19, 0.377 | -0.05, 0.984 |
| VentAttn ↔ Def+Lim | -0.59, 0.663 | -0.04, 0.969 | 0.03, 0.961 | -0.67, 0.700 | -1.12, 0.472 | -0.02, 0.984 |
| Executive ↔ Def+Lim | 0.89, 0.663 | 1.41, 0.547 | 0.56, 0.640 | 2.40, 0.045* | -0.72, 0.592 | 0.31, 0.984 |
| Maintain | 4.92, 0.757 | -0.34, 0.969 | 1.20, 0.961 | 5.85, 0.823 | 8.02, 0.592 | 3.97, 0.984 |
| Values are β estimates and FDR-adjusted p-values. * pFDR < 0.05; ** pFDR < 0.01; *** pFDR < 0.001. P-values corrected using FDR. The energetic costs were extracted from the uniform model. | | | | | | |
